# Systematic Identification of UBE2C As a Prognostic Biomarker and Correlated with Immunosuppression and Invasiveness in Glioma

**DOI:** 10.1101/2024.02.21.581365

**Authors:** Hao Feng, Anhui Fu, Rong Yang, Fei Qiao

## Abstract

Glioma is one of the common tumors of the central nervous system, which presents difficulties in clinical diagnosis and treatment due to its characteristics of immunosuppression and cell invasion phenotypes. If the condition and prognosis of glioma can be predicted during the process of diagnosis and treatment, it will be more conducive to timely intervention or evaluation of glioma. Therefore, we still need to search for more valuable tumor markers. The differential/risk genes and enrichment analysis based on glioma samples (The Cancer Genome Atlas, TCGA). Target gene UBE2C were obtained by the expression correlation and differential expression analysis for the enrichment results. UBE2C were evaluated by clinical grading, survival prognosis and cell experiments. The correlation of UBE2C with immune invasion, immune checkpoint, network analysis and cell invasiveness of gliomas was analyzed by TCGA-glioma data and STRING, respectively. The results suggests that the high expression and risk of UBE2C in gliomas may be a factor that promotes malignant phenotype of tumor cells. The immune phenotype shows that IL6 and IL10 may be the key nodes affecting the immunosuppressive phenotype of glioma. Further, the tumor cells aggressive genes from the MMP family can be correlated with immunosuppressive phenotypes via UBE2C-IL6/IL10 axis, especially displayed by MMP2/MMP9. The UBE2C may systemic effects the malignant phenotype, immunosuppression and cell invasiveness of tumors systematically, which reflects UBE2C as a potential biomarker of glioma and therapeutic target for this tumor.

## Introduction

As the main type of central nervous system tumor, brain glioma is also one of the most prone to malignant progression of tumors, which seriously endangers people’s nervous system health [1–3]. Currently, although this disease can be treated with surgery, radiotherapy and chemotherapy, the tumor is rapidly progressive, which prone to metastasis and immune checkpoints suppression [2, 4]. For example, the glioblastoma with a high degree of malignancy contains the above adverse characteristics [5]. Therefore, it is necessary to seek or explore relatively effective markers to effectively predict and evaluate the prognosis of patients with this tumor. Based on the malignant progression and high-risk characteristics of glioma, clarifying the correlation between this biomarker and the above characteristics becomes an important indicator of effective evaluation and prognosis [6, 7], and it may even be used as a potential target for glioma intervention.

The malignant phenotype of gliomas usually includes a highly aggressive and immunosuppressive phenotype, which is often a difficulty in clinical diagnosis and treatment [8, 9]. Therefore, the search for valuable biomarkers has become an important task for the effective prediction and prognosis of the disease. Compared with normal tissues, high-risk genes in glioma samples may be important factors in the adaptive survival or malignancy process of tumor cells [10]. UBE2C with relatively high expression in tumors may become typical risk factors. Studies have reported that UBE2C may play a role in promoting cancer, such as brain cancer, breast cancer, cervical cancer, pancreatic cancer and liver cancer [11, 12]. Based on this characteristic and combined with the malignant phenotype of glioma, multifaceted or systematic analysis can reflect that the expression level of this gene and may be a key factor affecting tumor progression [6, 13]. Such as immunosuppressive and highly aggressive phenotypes in glioma may be related to this gene, and thus become the key to the progression of tumor malignancy [6, 14]. These aspects mainly manifested by the systematic association of this gene with immune infiltrating cells, immune checkpoints and invasion-related genes [15]. Therefore, evaluating the intrinsic and extrinsic risk of these genes for glioma from a systemic perspective may be beneficial for obtaining valuable biomarkers.

Although there has been some progress in the study of biomarkers for glioma, systematic research on the relationship between risk genes and immunosuppressive or invasive phenotypes is rarely reported. Based on the TCGA database and GEO-sourced clinical glioma sample data (including glioblastoma multiforme and low-grade glioma), this study demonstrated that UBE2C gene could be used as a biomarker for glioma via systemic analysis, screening and evaluation. This gene has established a significant correlation with immunoinfiltrated Th2 cells, which may influence immunosuppressive phenotypes and tumor cell invasion via IL6 and IL10. In summary, based on these results, it can be reflected that the high expression of UBE2C in glioma may serve as a relatively typical risk factor and may become a valuable prognostic marker.

## Materials and Methods

### Glioma sample

The glioma samples involved in this study were taken from the TCGA database (https://www.cancer.gov/ccg/research/genome-sequencing/tcga), which contained 174 Glioblastoma multiforme, 532 low-grade gliomas, and 5 adjacent controls. In addition, there is a glioma sequencing dataset (GSE12657) from the GEO (Gene Expression Omnibus) database (https://www.ncbi.nlm.nih.gov/geo/), which contains five adjacent controls and seven Glioblastoma patients.

### Cell Culture and Transfection

The HBE, LN229, U87MG, A172 and U251MG cells involved in this experiment were all purchased from ATCC cell bank (http://www.atcc.org). All the cell lines were cultured in medium: DMEM (90 %, Gibco) + FBS (10% Fetal calf serum, Gibco) + 1 % PS (Penicillin streptomycin combination, Gibco), 37℃, 5% CO2. Cell transfection that transfection agents used for PolyPlus (https://www.polyplus-sartorius.com/), siRNA from Geneparma company (https://www.genepharma.com/). Please refer to the instructions of transfection reagent for the procedure of transfection experiment.

### Wound Healing and Trans-Well Assays

Tumor cells were seeded into 12 - well plates and scratched with a 10 μL pipette tip after reaching confluency. The plates washed thrice with PBS, cultured in serum-free medium DMEM for 24 h, and photographed. Cell migration assays, 10^4^ cells were spread into the filter of a 24 - well plate and cultured in serum-free medium DMEM for 48 h. Filters were then fixed with a neutral formaldehyde solution (4%) and stained with crystal violet.

### Colony Formation Assays and Cell Proliferation

Colony formation assay, cells were plated at a density of 1000 cells/well into 12-well plates and cultured for 14 days, then fixed with neutral formaldehyde (4 %) and stained with crystal violet. Cell proliferation assay was by seeding 5 × 10^3^ cells/well into a 96-well plate. Proliferation activity (Absorption in OD450 nm) was measured over four days using the CCK-8 method as per the manufacturer’s instructions.

### Western Blot

Protein samples extracted from cells were used for Western blotting, following a standard method. Reagents and consumables used in this process included PVDF membranes (Millipore, USA); anti-UBE2C (Rabbit, ab252940) (Abcam, USA); anti-GAPDH (Rabbit, ab181602) (Abcam, USA); TBST buffer (Biosharp, China); Running buffer (Biosharp, China); Transmembrane buffer (Biosharp, China); and Substrate luminescent liquid (Biosharp, China), Goat anti-Rabbit HRP (H+L, ab6721) (Abcam, USA).

### RNA Isolation and Real-Time PCR

Total RNA from tumor cells was extracted using the Trizol reagent method following standard procedures. Reverse transcription of total RNA was performed using a two-step kit (TaKaRa, Japan), and real-time qPCR experiments were conducted using the SYBR fluorescent dye kit (TaKaRa, Japan). The primers used in this procedure included: UBE2C (F: 5’-CATCAGAACCAGCTCAACAGT −3’; R: 5’-GTTGCAGAGTAAGCT CCAGCA −3’); GAPDH (F: 5′-ACAACTTTGGTATCGTGGAAGG −3′; R: 5′-GCC ATCACGCCACAGTTTC −3′).

### Immunohistochemical (IHC)

The glioma IHC images involved in this study are all from The Human Protein Atlas database (https://www.proteinatlas.org/), which contains the expression of existing related proteins and the IHC results of tumor pathological tissues (all images contain the sample number). Our conclusion is based on these image data for analysis and demonstration.

### GlioVis analysis

GlioVis online analysis (Visualization Tools for Glioma Datasets, http://gliovis.bioinfo.cnio.es/), which is an important database that adopted for data visualization and analysis to explore glioma [16]. Meanwhile, the normalized gene expression, this database includes information on glioma molecular pathology and glioma subtypes, which are important tools for online analysis.

### KEGG and GO enrichment analysis

The enrichment analyses involved in this study included KEGG (Kyoto Encyclopedia of Genes and Genomes) and GO (Gene Ontology). RNA-sequencing expression (level 3) profiles and corresponding clinical information for glioma were downloaded from the TCGA dataset (https://portal.gdc.com). Using the limma package in the R software to study the differentially expressed mRNA. “Adjusted *P* < 0.05 and Log_2_FC (Fold Change) > 1 or Log_2_FC < −1” were defined as the threshold for the differential expression of mRNAs. In addition, online Metascape analysis (https://metascape.org/gp/index.html#/main/step1) also serves as an important reference for enrichment analysis results [17].

### Gene Set Enrichment Analyses (GSEA)

Dataset GSE12657 was downloaded from the GEO database and then GSEA (http://software.broadinstitute.org/gsea/index.jsp) enrichment analysis was performed. The overall differential genes in this dataset were used as the data source for analysis without additional conditional screening. GSEA enrichments were estimated by normalized enrichment score (NES) [18]. The significance of the results was assessed with FDR < 0.25 level, *P* < 0.05, and FDR < 0.25 levels.

### Network system analysis

Online network analysis between multiple genes is performed using STRING (https://cn.stringdb.org/cgi/input?sessionId=bsNTRVpzqx25&input_page_show_search=on), and adjustments and layout optimization are made on this result. At the same time, the systematic analysis based on Gene Mania (https://genemania.org/) is used as a reference for the above results.

### Bioinformatics and statistical analysis

Gene expression differential analysis, clinical grading significance, survival prognosis analysis (K-M survival curve), immune infiltration analysis, gene expression correlation analysis, Cox risk regression analysis (Univariate and multivariate regression analysis) and nomogram analysis (Calibration curves) in gliomas were all implemented by R v4.0.3 software package. All the analysis results were represented by Spearman as the correlation coefficient (*R*), and *P* < 0.05 was the significant result.

## Results

### Function of Differentially Expressed Genes and Risk Genes in Glioma

Identifying the relatively typical biological processes in gliomas is the basis for seeking markers. Therefore, we first conducted differential gene enrichment analysis (KEGG and GO) based on glioma (containing the glioblastoma multiforme and low-grade glioma) sample data from TCGA database. Based on significantly upregulated genes, the enrichment results of both KEGG and GO reflected entries related to cell cycle regulation. We combined with the characteristics of tumor cell cycle, it was suggested that this process is the basis of malignant phenotype or progression of tumor cells. (**Figure 1A, B, where the red arrow is pointing**). Meanwhile, there were no typical characteristic items in the significantly down-regulated gene enrichment results (**Supplementary Figure 1A, B**). To further explore the commonality between the function of risk genes and up-regulated genes, 8344 risk genes (HR > 1, *P* < 0.05) were obtained by analyzing TCGA-glioma data, and they were compared with 1091 up-regulated genes (Log_2_FC > 1, *P* < 0.05) for Venn diagram analysis. The results showed that there were 679 genes with common characteristics (**Figure 1C**). Enrichment analysis of these genes revealed four biological processes associated with the cell cycle in the top 5 entries (**Figure 1D, where the red arrow is pointing**). These results suggest that upregulated risk genes in gliomas may also be involved in cell cycle processes.

**Figure 1.**
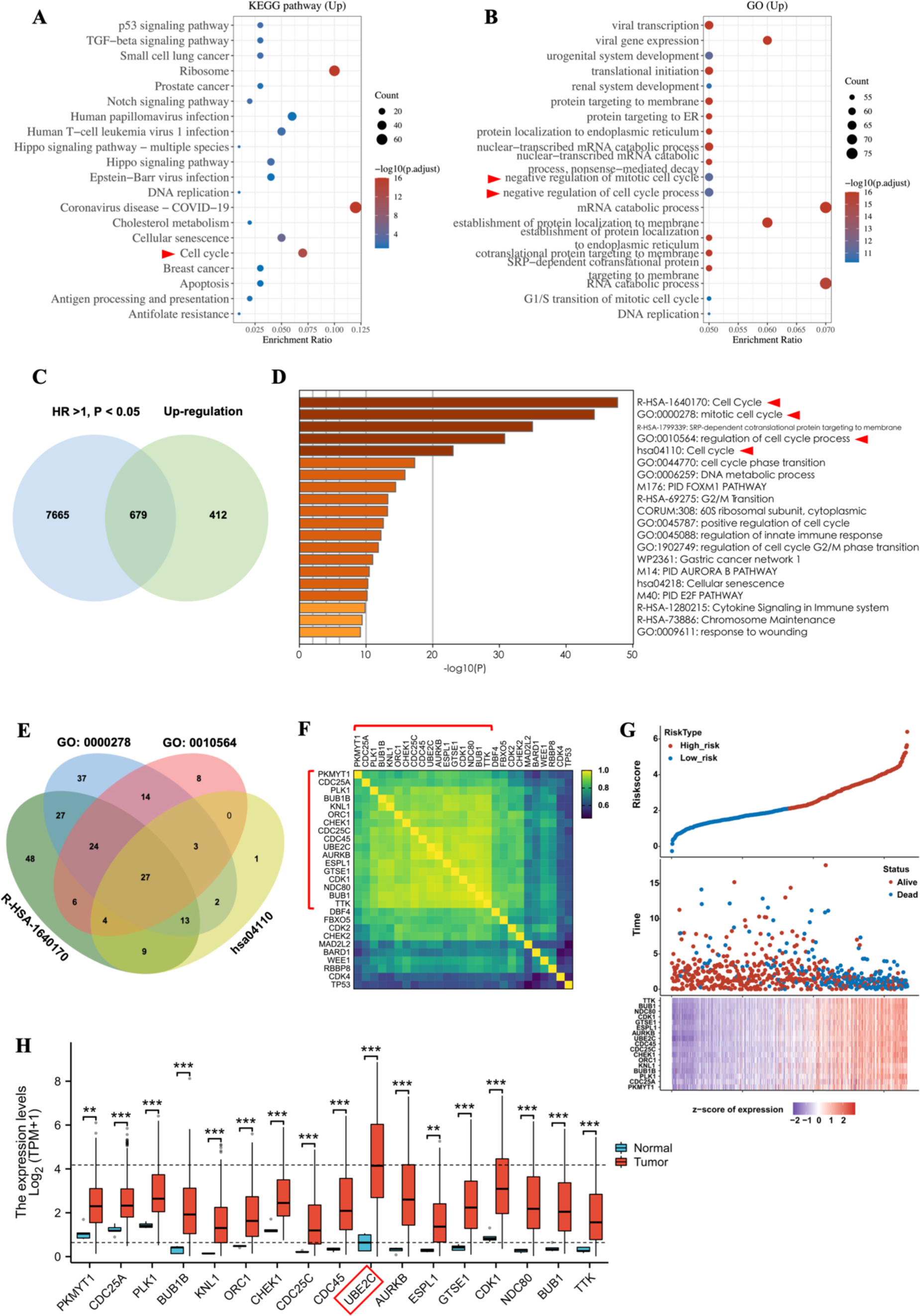
Screening of target genes in gliomas based on TCGA. **(A, B)** KEGG and GO enrichment analysis of differential genes (Upregulation, Log_2_FC > 1, *P* < 0.05), respectively; **(C)** Represents Venn diagram analysis of risk genes (HR > 1, *P* < 0.05) and up-regulated genes (Log_2_FC > 1, *P* < 0.05); **(D)** Enrichment analysis of 679 common genes from (C); **(E)** Venn diagram analysis of genes from the four biological processes in (D) (where the red arrow is pointing); **(F)** Analysis of expression associations among 27 genes from (E); **(G)** Co-expression and risk analysis of 17 genes from (F); **(H)** Differential expression analysis of 17 genes (The red box indicates that UBE2C has the highest differential expression). \*\**P* < 0.01, \*\*\**P* < 0.001.

Based on the above enrichment results, we conducted Venn diagram analysis for the genes from these four items, which found that 27 genes were the common part (**Figure 1E**). Correlation analysis of these genes in glioma showed that 17 genes were strongly correlated with each other (*R* > 0.8, *P* < 0.05), and all showed high risk at high expression levels (**Figure 1F, G**). These results confirmed the high-risk characteristics for the selected genes. Meanwhile, Cox regression analysis showed that 17 genes were significant risk factors in univariate analysis, and the multivariate prognostic analysis shows that CHEK1, UBE2C and BUB1 were independent prognostic risk factors (**Supplementary Figure 1C, D,** HR > 1, *P* < 0.05). In addition, according to the expression level of 17 genes in glioma, UBE2C had the most significant upregulation (**Figure 1H**, \*\*\**P* < 0.001). In summary, UBE2C may be a risk factor and significantly upregulated in gliomas, which may be the internal cause of malignant phenotype of tumors.

### Clinical Classification and Prognostic Significance of UBE2C in Glioma

UBE2C was selected as a typical representative gene. To further verify its clinical significance in glioma, these samples from the TCGA database were clinically classified (**Supplementary Table 1**). The results showed that the expression level of UBE2C was significantly up regulated with the progression of the disease in both WHO classification (G2, G3, G4) and primary therapy outcome (CR, Complete response; PR, Partial Response; SD, Stable disease; PD, Progressive disease) (**Figure 2A, B,** \*\**P* < 0.01, \*\*\**P* < 0.001). Among the pathological grades of glioma, the malignant degree of glioblastoma was the highest [5, 19], and the expression level of UBE2C was significantly higher than that of the other three types (**Figure 2C**, \*\*\**P* < 0.001). In terms of age and survival status, glioma patients over 60 years of age and those who had died showed significant UBE2C upregulation (**Figure 2D, E,** \*\*\**P* < 0.001). These features can also be demonstrated by the results of Cox regression analysis of clinical grading (**Supplementary Table 2**). These results suggest that the expression level of UBE2C is significantly different in the clinical grades of glioma, which can significantly affect the prognosis of these clinical grades.

**Figure 2.**
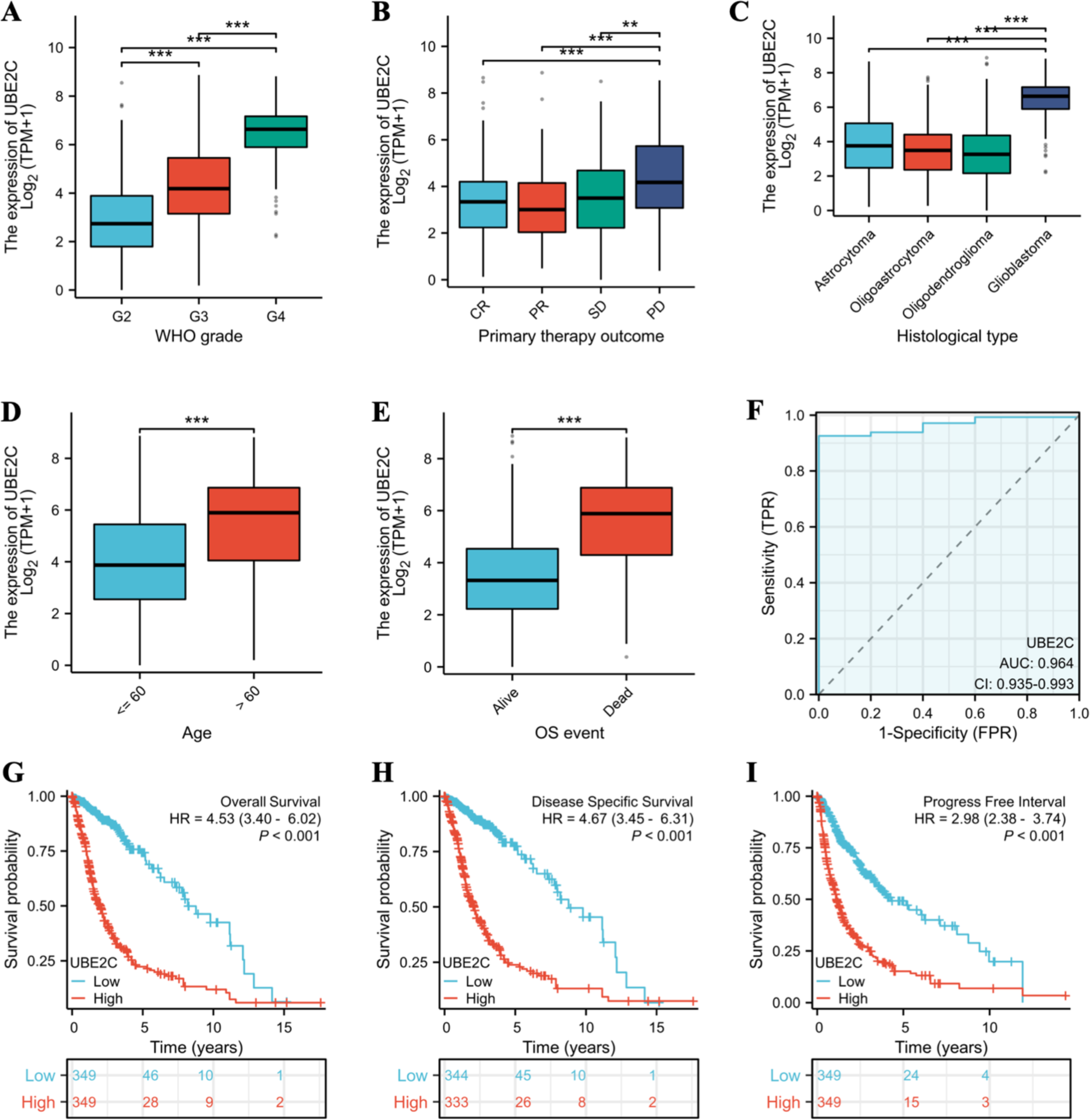
Clinical grading and prognostic analysis of UBE2C gene in TCGA-glioma samples. **(A)** The clinical grading based on WHO; **(B)** Represents the primary therapy outcome of glioma patients; **(C)** Indicates the histological type of glioma samples; **(D)** Represents the age division of glioma patients; **(E)** Represents the survival status of glioma patients; **(F)** Represents ROC curve analysis of glioma samples; **(G-I)** Respectively represent survival prognosis analysis based on TCGA-glioma with OS, DSS, PFI, respectively. \*\**P* < 0.01, \*\*\**P* < 0.001.

In addition, the ROC Curve (Receiver Operating Characteristic Curve) analysis of UBE2C in glioma samples showed that the AUC (Area Under ROC Curve, CI: 0.935 - 0.993) value was 0.964. This result indicates that UBE2C gene has good diagnostic value for glioma (**Figure 2F**). To assess the prognostic value of UBE2C in glioma, survival curves (K-M) with OS (Overall Survival), DSS (Disease Specific Survival), and PFI (Progress Free Interval) were performed on these clinical samples, respectively. The results showed that the three types of K-M curves showed that the survival rate of the group with high expression of UBE2C was relatively low, that is, the expression level of this gene was significantly negatively correlated with the survival rate of patients (**Figure 2G-I**). In conclusion, the differences in the expression of UBE2C in clinical classification and survival prognosis can reflect the related characteristics of disease progression, and thus show certain clinical diagnosis and prognostic value.

### The Intrinsic Effect of UBE2C in Glioma

Based on the clinical significance of UBE2C in gliomas, we need to further explore the internal influence of this gene in gliomas, which can reflect the value of UBE2C as a marker of endogenous adverse factors. We need to further explore the major biological processes involved in UBE2C by enrichment analyzing the genes that were significantly positively correlated with UBE2C (*R* > 0.6, adj. *P* < 0.05). The results show that the TOP 20 items presented mainly include two biological processes, which are cell cycle regulation (red arrow) and DNA behavior (blue arrow) (**Figure 3A**). The results of corresponding network and correlation analysis also reflect a fact that the cell cycle and DNA behavior are two major units, which is a close bond between them (**Figure 3B, C,** red and blue dashed line ellipses). GSEA analysis of differential genes based on TCGA verified the significance of cell cycle and DNA behavior, respectively (NES > 1, adj. *P* < 0.05, FDR q < 0.25) (**Figure 3D, E**). The analysis results of the GEO-sourced dataset (GSE12657) can also fully verify this conclusion (**Supplementary Figure 2A, B,** NES > 1, adj. *P* < 0.05, FDR q < 0.25). Therefore, the above results indicate that UBE2C may satisfy the adaptive survival of tumor cells in glioma mainly via the regulation of cell cycle and DNA behavior.

**Figure 3.**
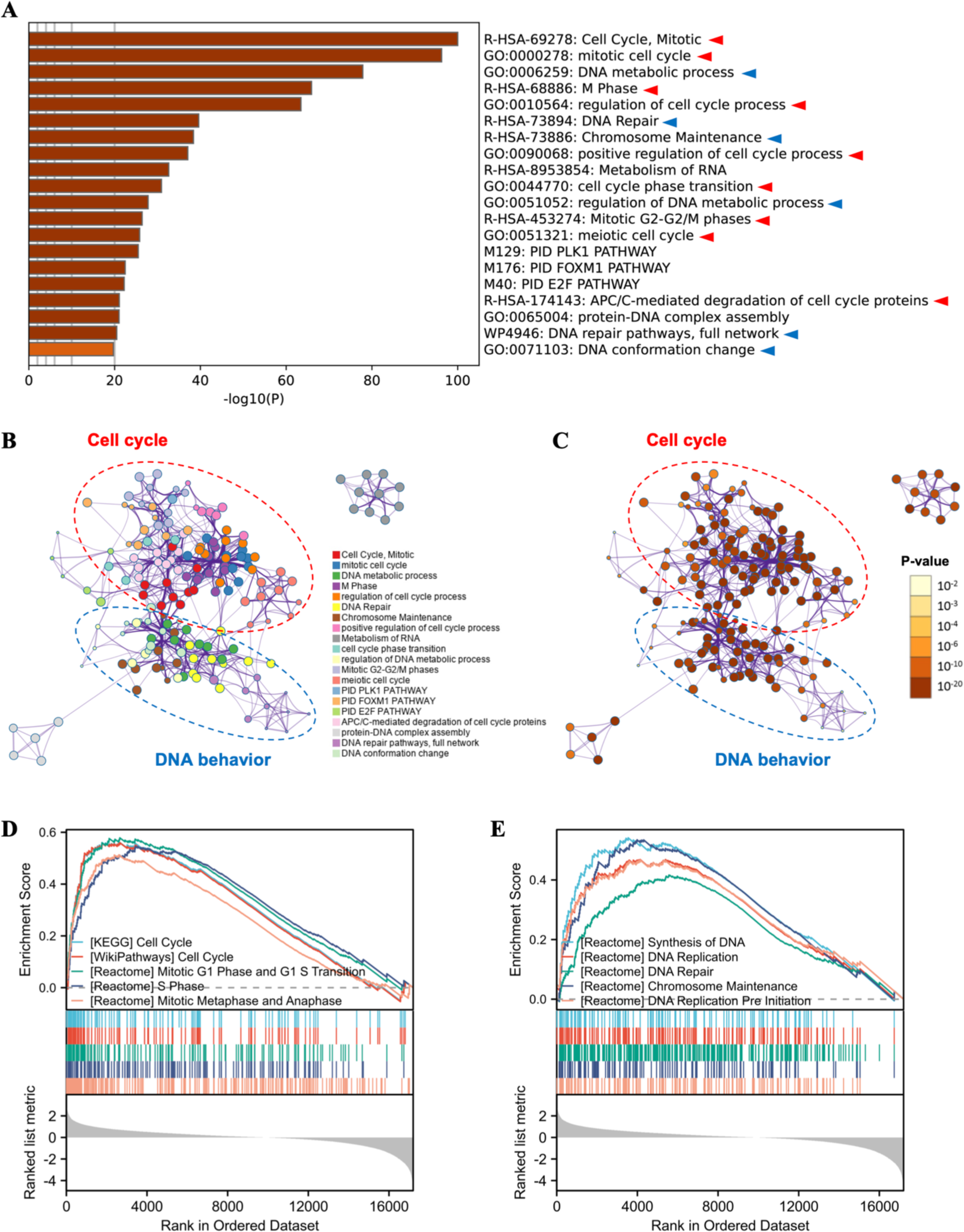
Enrichment analysis of genes positively related to UBE2C. **(A)** Represents enrichment analysis based on UBE2C positively related genes (red arrow represents cell cycle related processes, blue arrow represents DNA behavior related processes). **(B, C)** Represents the network analysis based on the results of the enrichment analysis in (A); **(D, E)** Represents a differential gene GSEA enrichment assay validation based on TCGA-glioma, which are cell cycle and DNA behavior processes, respectively. NES > 1, adj. *P* < 0.05, FDR q < 0.25.

Summarizing the above results, it can be inferred that UBE2C gene may be related to malignant proliferation of glioma cells. To verify this association, we separately display the IHC (Immunohistochemistry) images from The Human Protein Atlas (THPA) database. Since Ki67, PCNA and MCM7 are common markers reflecting tumor cell proliferation [20], they are related to the replication behavior of nuclear DNA. Therefore, according to the IHC of these three proliferative markers, the nuclear coloring degree of tumor region is deeper than adjacent region (**Figure 4B-D**). Meanwhile, the staining trend of UBE2C cells in tumor region was consistent with these three proliferative markers (**Figure 4A**). In addition, the scatter plot of correlation analysis shows that UBE2C had a significant positive correlation with Ki67, PCNA and MCM7, respectively (**Figure 4E-G**). In summary, the expression level of UBE2C is related to the proliferation of tumor cells, and it may be an internal factor affecting the adaptive survival of tumor cells.

**Figure 4.**
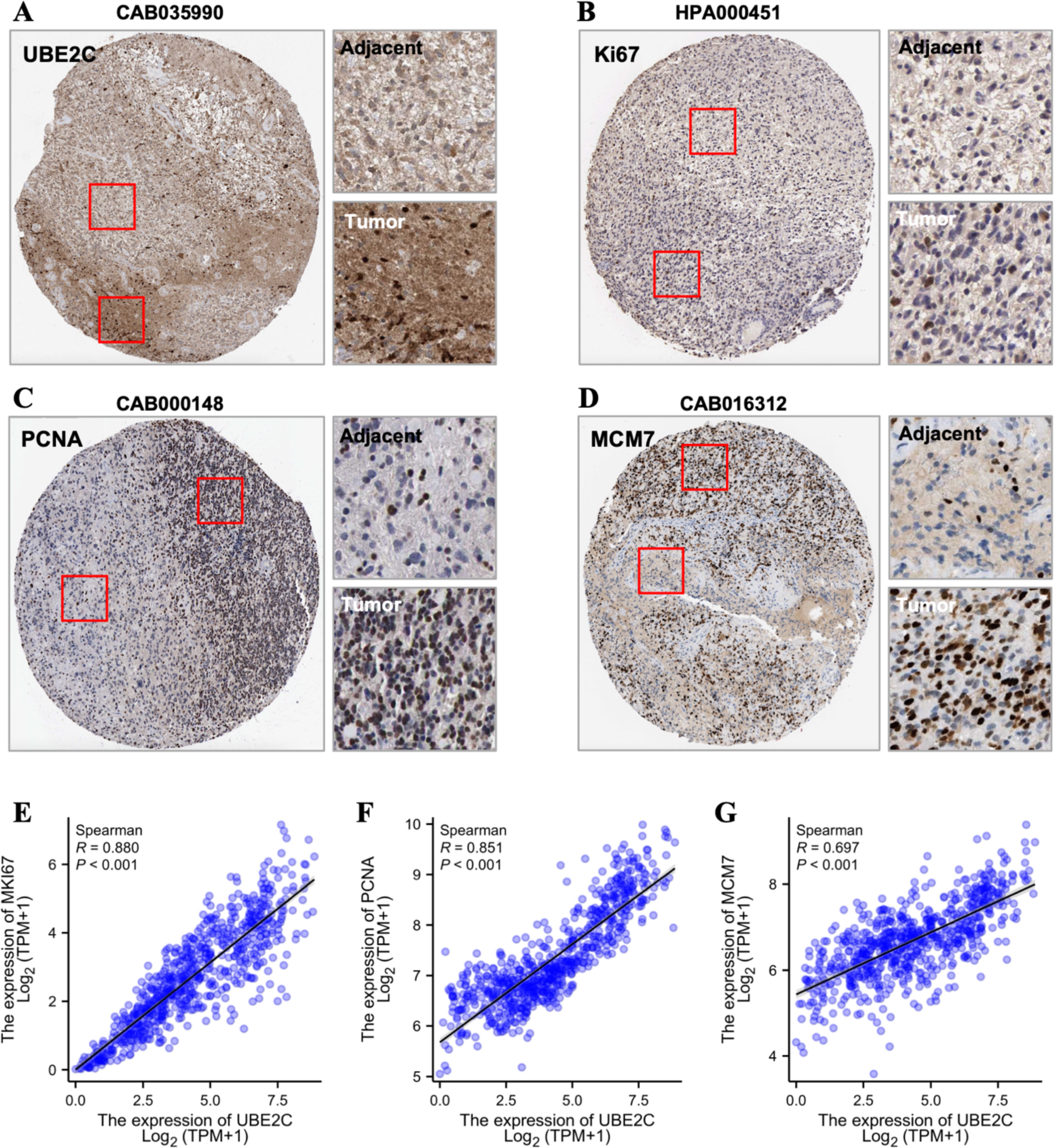
Expression levels of UBE2C and cell proliferation antigen in gliomas. **(A-D)** Represents IHC images of UBE2C, Ki67, PCNA and MCM7 from The Human Protein Atlas database, respectively; **(E-G)** Represents the scatter plot of expression correlation between UBE2C and Ki67, PCNA and MCM7, respectively; **(H)** Expression levels of UBE2C mRNA in 48 brain cancer cell lines from the CCLE database; **(I)** The expression level of UBE2C mRNA in 5 cell lines was detected by qPCR; **(J)** The expression level of protein UBE2C in 5 cell lines was detected by Western blot.

To verify the expression of UBE2C in glioma cell lines, our analysis based on the mRNA data collected in the CCLE database (Cancer Cell Line Encyclopedia), which showed that there was little difference in the expression of UBE2C in 48 brain cancer cell lines (**Figure 4H**). Meanwhile, the differences between the common glioma cell lines LN229 and U251MG were relatively obvious (**Figure 4H**, where the red arrow is pointing). The expression of UBE2C in 4 glioma cell lines (LN229, U87MG, A172 and U251MG) was detected by qPCR and Western blot, respectively. The results showed that the mRNA expression level of UBEC in these four cell lines was significantly higher than that of normal glioma cells (HEB), but there was no difference among the four cells (**Figure 4I**, \*\**P* < 0.01, \*\*\**P* < 0.001). In addition, the expression of UBE2C protein in the four types of glioma cells was similar to mRNA levels (**Figure 4J**, \**P* < 0.05, \*\**P* < 0.01, \*\*\**P* < 0.001). In conclusion, the relatively high expression level of UBE2C in glioma cell lines was consistent with TCGA data and IHC results.

### Effect of UBE2C Expression on Malignant Phenotype for Glioma Cell

Based on the difference of UBE2C expression in glioma (tissue and cellular level) and its relevance to the clinical prognosis of this tumor, we need to further verify the effects of UBE2C on the malignant phenotype for the glioma at the cellular level. UBE2C was highly expressed in glioma cell lines LN229 and U251MG (**Figure 4H, J**). Therefore, we achieved knockdown of UBE2C protein expression via transfecting these two cells with siRNA. The results showed that siRNA3 had the most significant knockdown effect on these two types of cells (**Figure 5A, B,** \**P* < 0.05, \*\**P* < 0.01). Since the knockdown effects of siRNA3 above, cell scratch experiments showed that the knockdown of UBE2C in both LN229 and U251MG cells could significantly inhibit the healing ability of these two cells (**Figure 5C, D,** \*\**P* < 0.01, \*\*\**P* < 0.001). In addition, when UBE2C is knocked down in LN229 and U251MG cells, it can significantly inhibit the migration ability of these two cells (**Figure 5E, F,** \*\**P* < 0.01). These results indicate that the expression level of UBE2C is positively correlated with the migration ability of tumor cell lines LN229 and U251MG.

**Figure 5.**
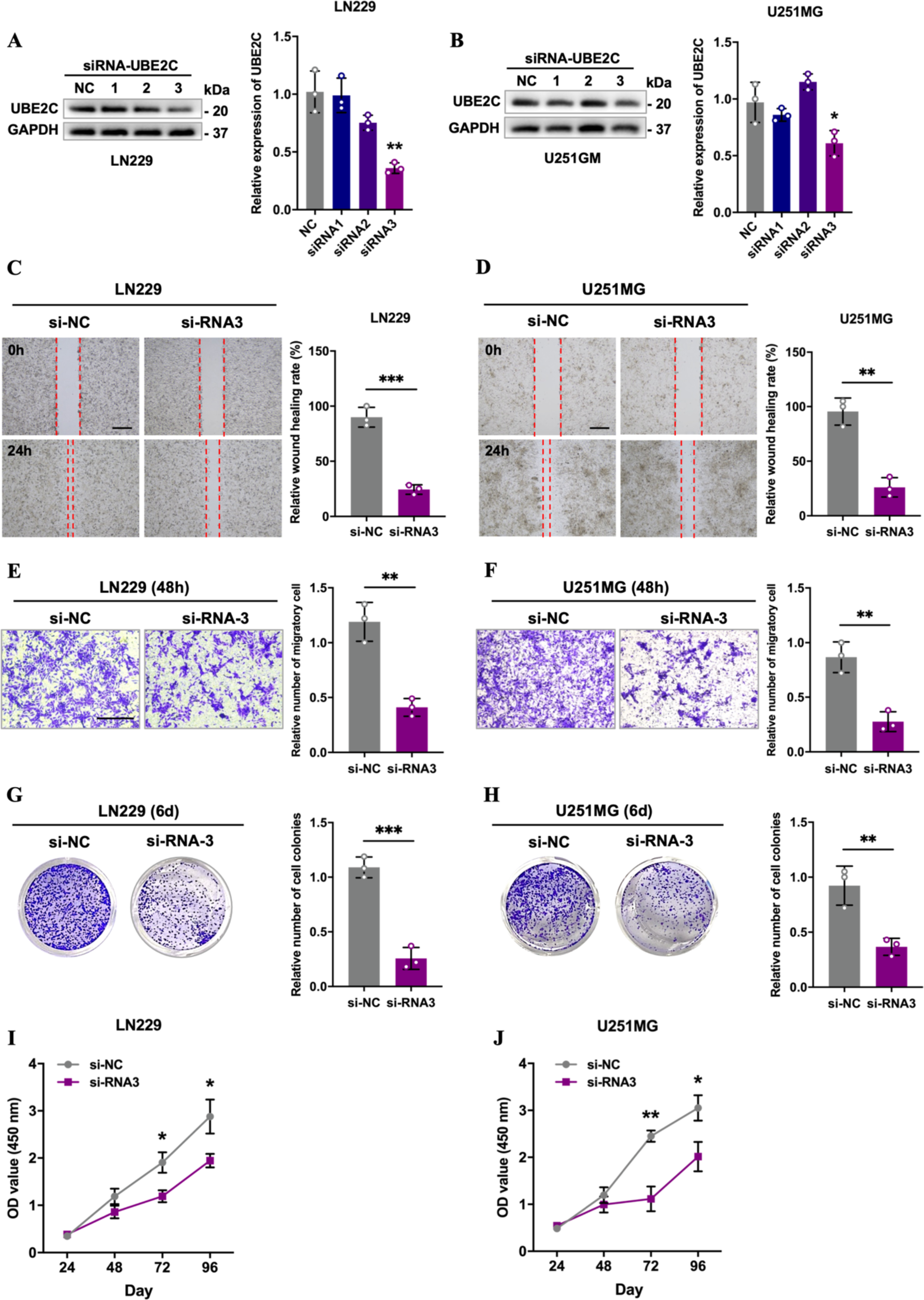
Effect of UBE2C Expression on Malignant Phenotype for Glioma Cell. **(A, B)** The knockdown effect of three siRNA on LN229 and U251MG cells was evaluated by Western blot, respectively; **(C, D)** The healing ability of UBE2C - siRNA3 for LN229 and U251MG cells was evaluated by cell scratch assay, respectively; **(E, F)** The migration ability of UBE2C - siRNA3 for LN229 and U251MG cells was evaluated by trans-well assay, respectively; **(G, H)** The proliferation ability of UBE2C - siRNA3 for LN229 and U251MG cells was evaluated by cell clonal formation assay, respectively; **(I, J)** The proliferation ability of UBE2C - siRNA3 for LN229 and U251MG cells was evaluated by CCK8 assay (OD450), respectively. \**P* < 0.05, \*\**P* < 0.01, \*\*\**P* < 0.001.

In addition, the cell cloning assay of LN229 and U251MG showed that when UBE2C was knocked down, it could significantly inhibit the ability of clonal formation for the two tumor cells (**Figure 5G, H,** \*\**P* < 0.01, \*\*\**P* < 0.001). Meanwhile, the cell proliferation curve assay (OD450, CCK8) showed that the knockdown of UBE2C could significantly inhibit the proliferation ability of these two types of cells (**Figure 5I, J,** \**P* < 0.05, \*\**P* < 0.01). Therefore, these results indicate that the expression level of UBE2C can significantly affect the malignant phenotype of LN229 and U251MG cells.

### The External Effects of UBE2C in Glioma

As an intrinsic risk factor affecting the progression of glioma, UBE2C may be involved in the regulation of cell cycle to positively link the proliferative activity of tumor cells. However, we also need to understand its correlation with external factors. Based on TCGA data, we analyzed the correlation between UBE2C and 24 types of immune cell infiltration in glioma. The results showed that the expression level of UBE2C was most significantly correlated with the immune infiltration of Th2 cells (**Figure 6A**, *R* = 0.871, \*\*\**P* < 0.001). Therefore, we are concerned that this feature may be the focus as UBE2C is linked to external factors of the tumor. According to the secretory phenotype of Th2 cells [21, 22] and the trend of UBE2C expression, and combined with their correlation analysis, we can see that IL6 (R = 0.396, *P* < 0.01) and IL10 (R = 0.338, *P* < 0.01) are the most significant (**Figure 6B, C**). In conclusion, UBE2C may be closely related to the immune infiltration of Th2 cells.

**Figure 6.**
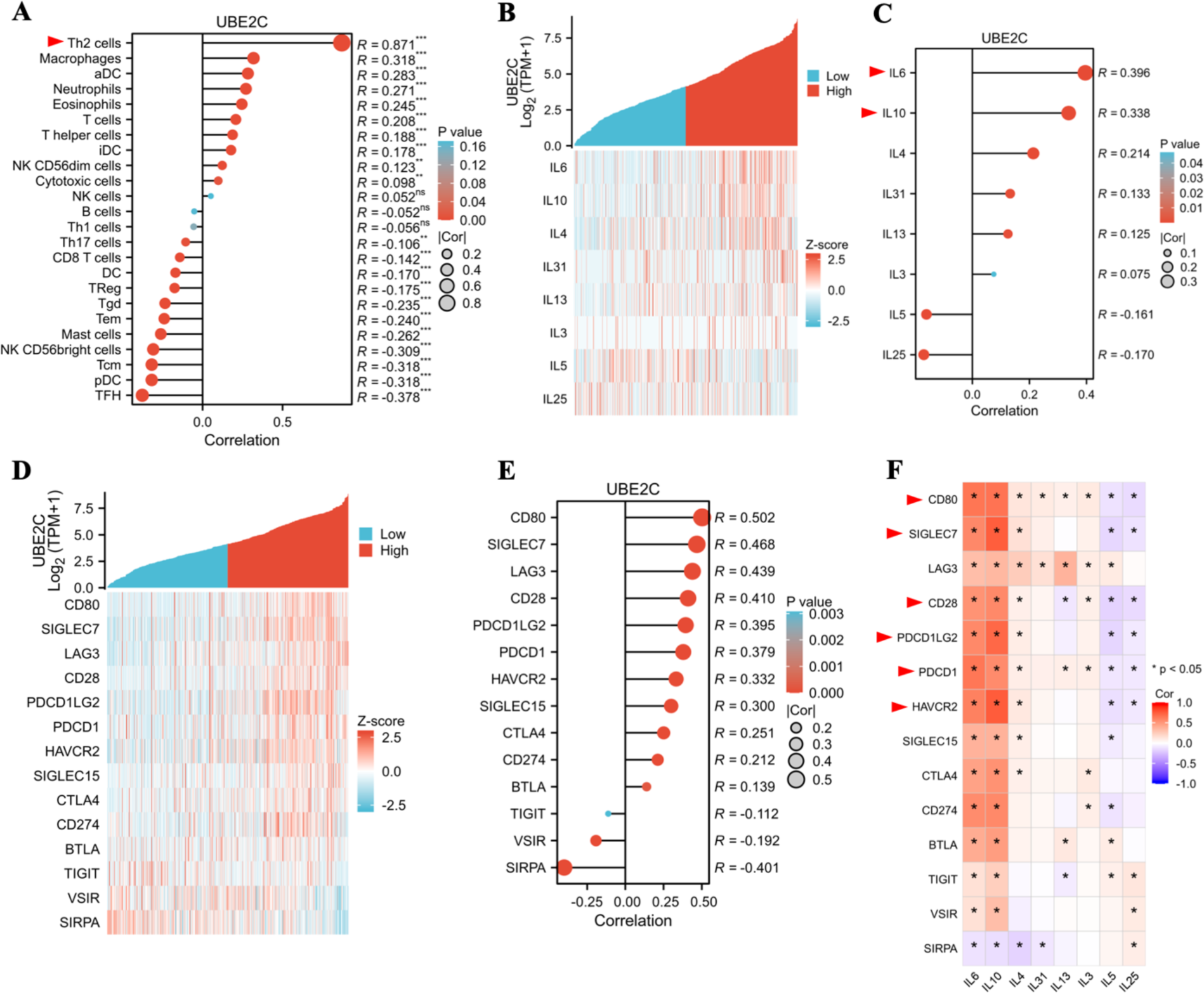
Correlation analysis of UBE2C with immune cell infiltration and immune checkpoint based on TCGA-glioma. **(A)** Correlation analysis between UBE2C and immune cell infiltration; **(B)** Co-expression analysis between UBE2C and secretory phenotypes from Th2 cells; **(C)** correlation analysis between UBE2C and secretory phenotype of Th2 cells; **(D)** Analysis of co-expression between UBE2C and immune checkpoints; **(E)** Correlation analysis between UBE2C and immune checkpoints; **(F)** Correlation analysis between immune checkpoint and Th2 cell secretion phenotype.

It has been reported that IL6 or IL10 can induce immunosuppressive phenotypes in the tumor microenvironment, which leads to the immune escape mechanism of tumor cells [23]. Therefore, for malignant progression and poor prognosis of glioma, the correlation between UBE2C and external adverse factors may reflect the essence behind this phenomenon. We analyzed the correlation between the expression of UBE2C and 14 immune checkpoints in glioma, and the results showed that eight immune checkpoints (CD80, SIGLEC7, LAG3, CD28, PDCD1LG2, PDCD1, HAVCR2, SIGLEC15) were significantly positively correlated with UBE2C (**Figure 6D, E,** *R* > 0.3, *P* < 0.001). Based on the correlation analysis of Th2 cells secretion phenotype and immune checkpoints, we found that IL6 and IL10 were most significantly correlated with various immune checkpoints, among which six immune checkpoints were more typical (where the red arrow is pointing) (**Figure 6F**, *R* > 0.5, \**P* < 0.05). These results suggest that secretory phenotypes IL6 and IL10 from Th2 cells may be important factors affecting immunosuppressive phenotypes in gliomas.

Based on the systematic correlation characteristics of UBE2C - IL6/IL10 - immune checkpoint axis, we further analyzed the prognostic value of glioma. The results showed that the expression levels of 13 genes, such as IL6/IL10 and immune checkpoint, were significantly negatively correlated with the survival rate of patients (**Figure 7A**). Meanwhile, IHC images from the THPA database displayed relatively typical expressions of six proteins, which indicated that IL6, IL10, PDCD1LG2, PDCD1, HAVCR2 and CD80 were upregulated in glioma tissues, respectively (**Figure 7B-G**). Taking together, the expression levels of UBE2C-related immune infiltrating Th2 cells and immune checkpoints can significantly affect the survival and prognosis of glioma, which suggesting that UBE2C may be an external risk factor to predict tumor progression.

**Figure 7.**
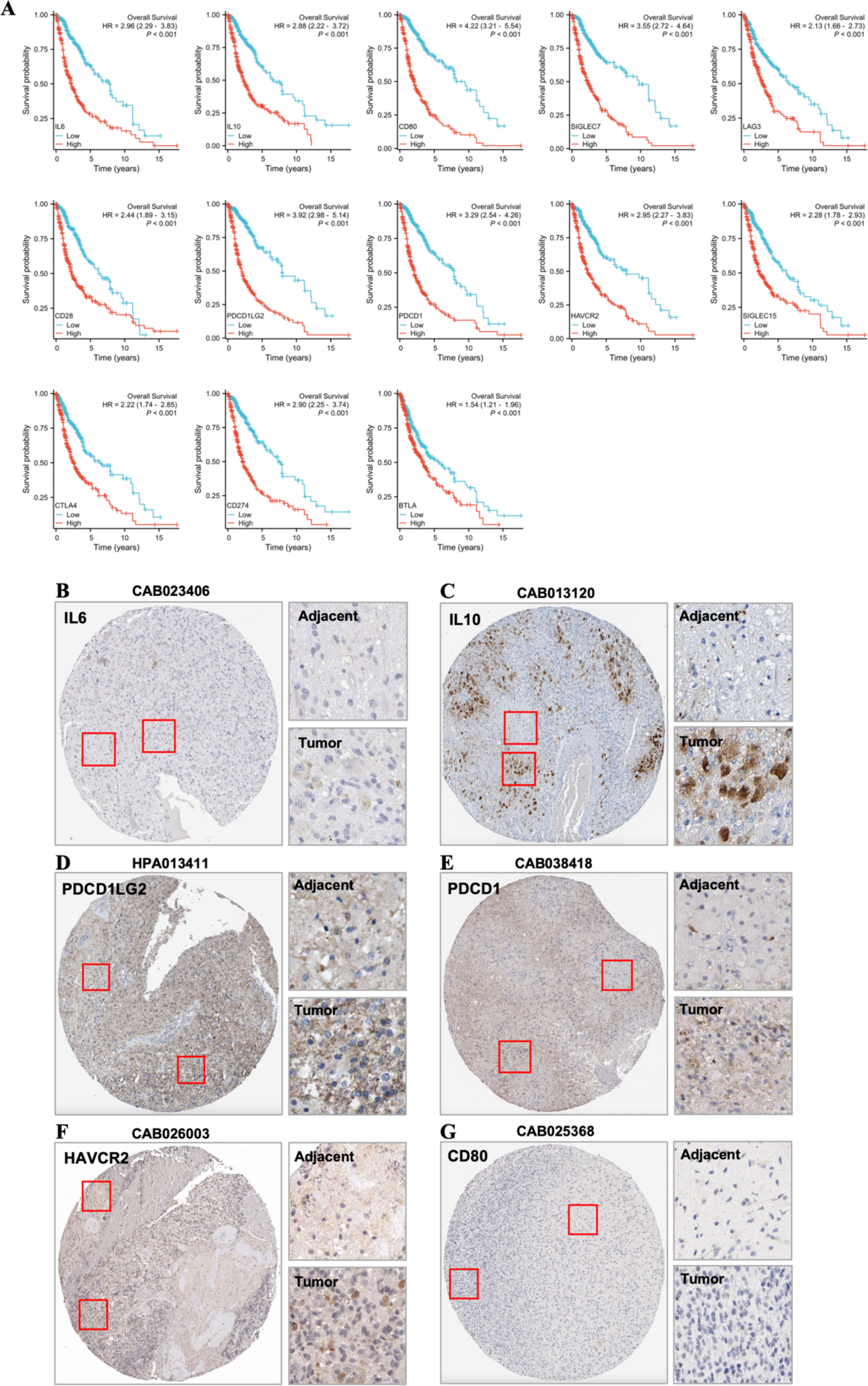
Expression and prognostic survival analysis based on immune checkpoint and typical Th2 cell secretion phenotype. **(A)** Prognostic survival analysis based on immune checkpoint and typical Th2 cell secretion phenotype; **(B-G)** Represents IHC images of IL6, IL10, PDCD1LG2, PDCD1, HAVCR2 and CD80 from THPA database, respectively.

### Correction of UBE2C-Related Risk Factors with Cell Invasion in Glioma

Since the high aggressiveness of glioma cells has been a major difficulty in the treatment of this disease [24, 25], we should also consider its association with cell aggressiveness phenotype when exploring the internal and external correlation of UBE2C. It has been reported that the invasiveness of tumor cells is correlated with soluble immune checkpoints, which reflects that immune checkpoints in the immune microenvironment can significantly affects the invasive phenotype of tumor cells [26]. Therefore, we performed network and correlation analyses of between MMPs (Matrix Metalloproteinase) family genes and immune checkpoints, which due to the MMP family plays an important role in the invasion of tumor cells [27]. Based on these backgrounds and analysis, we found that UBE2C-related risk genes (IL6, IL10 and immune checkpoints) were significantly positively correlated with MMP1, −2, −3, −7, - 8, −9, −10, −11, −12, −13, −14 and −19, respectively (**Figure 8A**, \**P* < 0.05). In addition, online analysis by STRING showed that UBE2C, IL6, IL10 and MMP9 acted as a “linker” that between immune checkpoints and the MMP family (**Figure 8B**). Subsequently, after verifying the correlation between seven MMPs (The most typical seven MMPs, which including MMP1, −2, −7, −9, −11, −14, −19) and immune infiltration in glioma, it was found that MMP2, MMP9 and MMP11 had the most significant correlation with Th2 cells in glioma (*R* > 0.5, *P* < 0.001) (**Figure 8C-E, Supplementary Figure 3A-D**). In conclusion, UBE2C, IL6 and IL10 may be the link between immune checkpoint and MMP family. This result also confirmed that the immune invasive phenotype (IL6, IL10 and immune checkpoints) of Th2 may be related to the aggressiveness of tumor cells.

**Figure 8.**
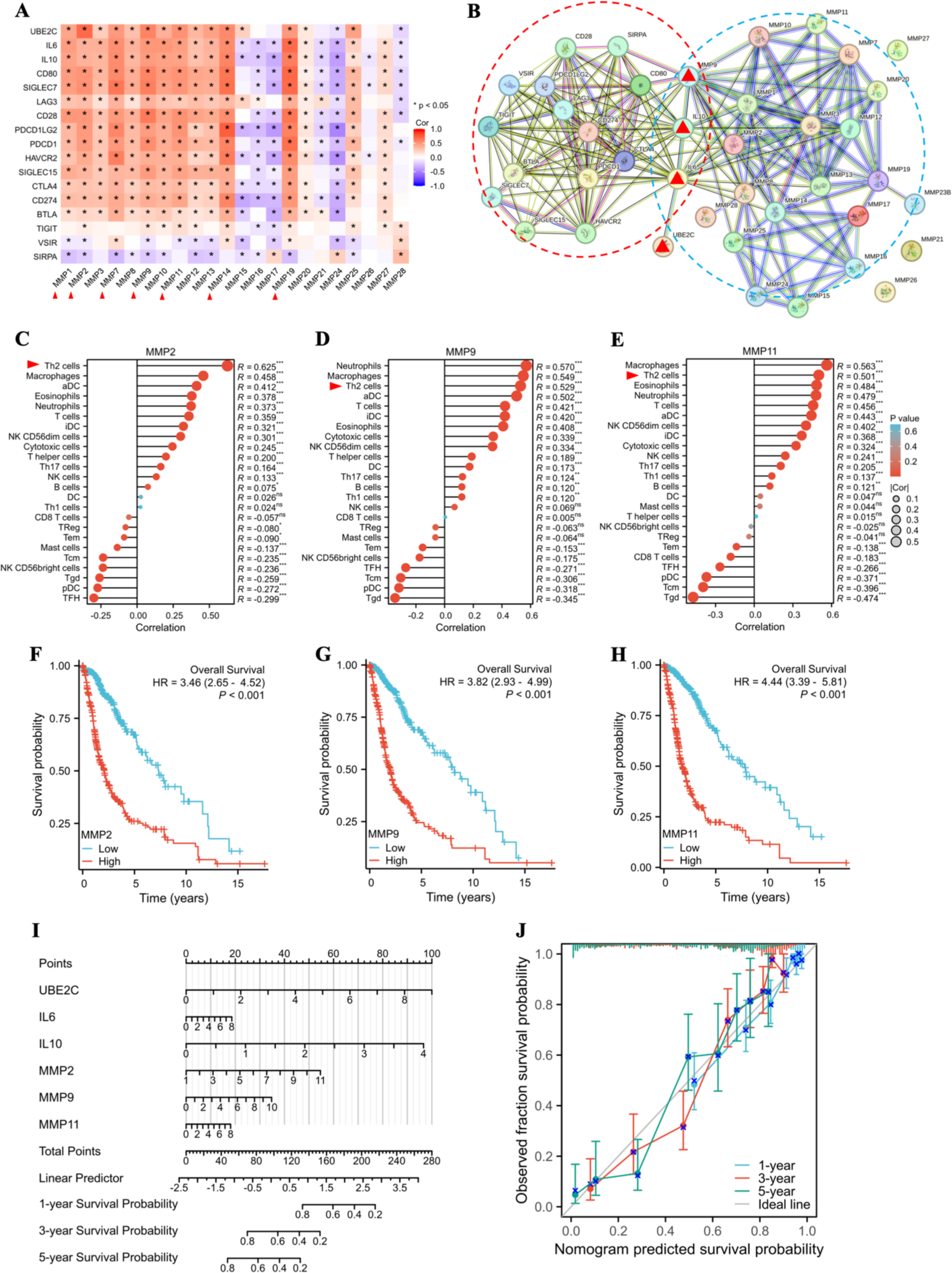
Correction of UBE2C-related risk factors with cell invasion in glioma. **(A)** Correlation analysis between immune checkpoint and MMP family expression; **(B)** Analysis of network correlation between immune checkpoint and MMP family by STRING online; **(C-E)** Analysis of the correlation between MMP2, MMP9 and MMP11 and immune cell infiltration, respectively; **(F-H)** Analysis of survival prognosis of MMP2, MMP9 and MMP11 in glioma patients, respectively; **(I, J)** The prognostic nomogram and calibration curve of UBE2C, IL6, IL10, MMP2, MMP9, and MMP11 in glioma patients, respectively.

Based on these findings, we need to further clarify the prognosis of these seven MMPs for the glioma patients. K-M survival analysis (OS) showed that the expression levels of seven MMPs were significantly negatively correlated with the survival rate of glioma patients, and the MMP2, MMP9 and MMP11 were more typical (**Figure 8F-H, Supplementary Figure 3E-H**). IHC images from the THPA database showed that MMP2 and MMP14 expression differences were most significant in gliomas (MMP1 and MMP19 data were not included) (**Supplementary Figure 4A-E**). Finally, multivariate Cox regression models of UBE2C, IL6, IL10, MMP2, MMP9 and MMP11 were analyzed by nomogram to predict 1-, 3-, and 5-year survival probabilities. The results showed that the contribution value of UBE2C, IL10 and MMP2 was the largest (total score ratio: 250/280 = 89.3%), which may be used as a more accurate multi-factor prediction model for glioma patients, and the prediction model was consistent with the conclusion, that is, UBE2C was correlated with immune invasion and cell invasion (**Figure 8I**). Meanwhile, the calibration curve verifies that the 1-, 3-, and 5-year survival probability curves fit well with the ideal line based on the same conditions, which indicates that the multi-factor prediction model is reliable (**Figure 8J**). In conclusion, among the risk factors for glioma, UBE2C may be systematically associated with IL10 and MMP2, which become a better predictive model. Therefore, UBE2C may be an ideal biomarker for the prognosis of glial patients.

## Discussion

Glioma is a major tumor that seriously threatens the health of the nervous system, which includes a variety of diseases with different degrees of malignancy [1, 28, 29]. At present, the difficulty of clinical treatment of glioma is the malignant progression and poor prognosis from the tumor [1, 30]. However, there has been some progress in the study of gliomas, especially in the molecular mechanisms of the tumor microenvironment and cell invasion [31–33]. However, the search for effective diagnosis and treatment of clinical tumors and prognosis assessment is still one of the tasks that need to be solved [2, 6]. The clinical application of biomarkers for glioma is mainly reflected in pathological diagnosis, prediction and prognosis assessment, which is also based on the gene expression profile, mutant and the expression difference of risk genes between tumor cells and normal cells to screen [6, 34, 35]. The purpose of this study was to obtain candidate markers via the differential expression of risk genes in the overall sample of gliomas (including glioblastoma and low-grade gliomas) (**Figure 1, 2, Supplementary Figure 1**). It is characterized by the possibility of considering the range of gliomas with different degrees of malignancy, which should be an important angle to search for potential biomarkers of gliomas. Currently, non-coding RNA, circulating exosome factors and cerebrospinal fluid can be the scope of seeking glioma biomarkers [7, 34, 36–38]. However, this study suggests that in addition to circulatory source factor expression differences and prognostic analysis, systematic analysis and evaluation based on markers are also key to validate their reliability [39, 40], such as assessing the intrinsic and extrinsic association and risk of marker genes, which are also needed. Therefore, a relatively typical marker - UBE2C was selected based on systemic intrinsic factors such as differential expression genes, risk genes, cell cycle and DNA replication in glioma. However, in terms of the reliability of prognostic markers, validation of external associations is still necessary.

Studies have reported that UBE2C, as an E2 ubiquitin binding enzyme, is involved in the regulation of glioma autophagy, chemotherapy resistance, aggressiveness, and poor prognosis [41–43]. However, systematic analysis to evaluate this gene as a potential marker has not been reported. Based on this background, this study not only analyzed the prognostic effect of UBE2C in glioma from multiple perspectives (**Figure 1, 2**), but also found that this gene is related to cell cycle and DNA behavior (**Figure 3**, **4**). These biological processes may be the intrinsic reasons for glioma adaptive survival, and the correlation between UBE2C and malignant phenotypes of glioma cell also reflects this inference (**Figure 5**). In addition, since the immunosuppression and cell invasion phenotypes of glioma are difficult in clinical diagnosis and treatment [44, 45], the evaluation of the application value of biomarkers should also focus on these two aspects [46–48]. In this study, UBE2C was found to be significantly positively correlated with immune infiltration of Th2 cells in gliomas (**Figure 6A-C**). Therefore, according to the secretory phenotype of Th2 cells, it is speculated that Th2 cells may be an important factor leading to immunosuppressive phenotype in the glioma microenvironment, which has been reported in relevant studies [49, 50]. Our correlation analysis results confirmed the conjecture that IL6 and IL10 were significantly positively correlated with most immune checkpoints (**Figure 6F**). This conclusion confirms the extrinsic association between UBE2C and glioma, which may be a risk factor for the adaptive progression of tumor cells.

The aggressiveness of tumor cells is also one of the main malignant phenotypes of glioma, which may be regulated and influenced by the tumor microenvironment [51, 52]. It has been reported that immune checkpoints can affect the expression of MMPs in tumor cells, which promoting the aggressiveness of tumor cells [53, 54]. Therefore, exploring the correlation between UBE2C and MMP gene family is also an important aspect of the external correlation between UBE2C and glioma. In this study, the network or association analysis of between immune checkpoints and MMP families showed that IL6, IL10 and MMP9 correlated with UBE2C may be a “linker” of two extrinsic phenotypes, which is reflected that a significant correlation (**Figure 8A, B**). This feature highlights the potential value of UBE2C as a biomarker for glioma. In summary, we identified UBE2C as a potential prognostic biomarker for glioma, and these results demonstrated the internal or external correlation and effects of UBE2C in glioma, but this causal relationship need to be further verified and explored by experiments. Based on the prognostic value and correlation of UBE2C, it may systematically reveal the factors of glioma cells’ adaptive survival in the tumor microenvironment, and further reflect that UBE2C may be a potential target for the treatment of this disease.

## Declarations

### Competing interests

The authors declare that they have no competing interests.

### Author contributions

Study concept and design: HF and FQ; Acquisition of data: HF, AF and RY; Analysis and interpretation of data: HF, AF; Statistical analysis: HF; Drafting of the manuscript: HF and FQ; Manuscript check: HF, RY and FQ; Critical revision and final approval of the manuscript: FQ. All authors contributed to the article and approved the submitted version.

### Ethics Statement

The studies involving data and platform were reviewed and approved by The Institutional Research Ethics Committee of Nanchong Central Hospital.

### Consent for publication

All personal data and samples involved in this study have been obtained with their knowledge and permission for publication.

### Data Availability Statement

All datasets generated and analyzed during the current study are available from the corresponding authors on request.

### Funding

This study was not supported by any funding.

## Acknowledgements

We are especially grateful for the relevant personnel of the Nanchong Central Hospital/Department of Neurosurgery for their platform and help.

---

**Supplementary Figure 1. (A, B)** KEGG and GO enrichment analysis bubble maps based on differential genes (TCGA-glioma, down-regulation), respectively; **(C, D)** Represents Forest maps of univariate and multivariate Cox regression analyses based on 17 genes, respectively.

**Supplementary Figure 2. (A, B)** Represents the results of GSEA enrichment analysis based on dataset GSE12657, which includes biological processes of cell cycle and DNA behavior, respectively.

**Supplementary Figure 3. (A-D)** Indicates the correlation analysis of MMP1, MMP7, MMP14 and MMP19 with immune cell infiltration in TCGA-glioma, respectively; **(E-H)** Represents survival prognosis analysis for MMP1, MMP7, MMP14 and MMP19 in TCGA-glioma, respectively.

**Supplementary Figure 4. (A-E)** The expression level of MMP2, MMP7, MMP9, MMP11 and MMP14 in glioma tissues was displayed by IHC images from THPA database, respectively (MMP1, MMP4 and MMP19 MMP1, MMP4 and MMP19 are not included in the THPA database).

## Notes

### Competing Interest Statement

The authors have declared no competing interest.

## References

1. Lapointe S, Perry A, Butowski NA: Primary brain tumours in adults. Lancet 2018, 392:432–446.

2. Yang K, Wu Z, Zhang H, Zhang N, Wu W, Wang Z, Dai Z, Zhang X, Zhang L, Peng Y, et al: Glioma targeted therapy: insight into future of molecular approaches. Mol Cancer 2022, 21:39.

3. Li T, Li J, Chen Z, Zhang S, Li S, Wageh S, Al-Hartomy OA, Al-Sehemi AG, Xie Z, Kankala RK, Zhang H: Glioma diagnosis and therapy: Current challenges and nanomaterial-based solutions. J Control Release 2022, 352:338–370.

4. Rong L, Li N, Zhang Z: Emerging therapies for glioblastoma: current state and future directions. J Exp Clin Cancer Res 2022, 41:142.

5. Schaff LR, Mellinghoff IK: Glioblastoma and Other Primary Brain Malignancies in Adults: A Review. JAMA 2023, 329:574–587.

6. Sledzinska P, Bebyn MG, Furtak J, Kowalewski J, Lewandowska MA: Prognostic and Predictive Biomarkers in Gliomas. Int J Mol Sci 2021, 22.

7. Jones J, Nguyen H, Drummond K, Morokoff A: Circulating Biomarkers for Glioma: A Review. Neurosurgery 2021, 88:E221–E230.

8. Mirzaei R, Sarkar S, Yong VW: T Cell Exhaustion in Glioblastoma: Intricacies of Immune Checkpoints. Trends Immunol 2017, 38:104–115.

9. Habashy KJ, Mansour R, Moussalem C, Sawaya R, Massaad MJ: Challenges in glioblastoma immunotherapy: mechanisms of resistance and therapeutic approaches to overcome them. Br J Cancer 2022, 127:976–987.

10. Barthel L, Hadamitzky M, Dammann P, Schedlowski M, Sure U, Thakur BK, Hetze S: Glioma: molecular signature and crossroads with tumor microenvironment. Cancer Metastasis Rev 2022, 41:53–75.

11. Yuan L, Yang Z, Zhao J, Sun T, Hu C, Shen Z, Yu G: Pan-Cancer Bioinformatics Analysis of Gene UBE2C. Front Genet 2022, 13:893358.

12. Xie C, Powell C, Yao M, Wu J, Dong Q: Ubiquitin-conjugating enzyme E2C: a potential cancer biomarker. Int J Biochem Cell Biol 2014, 47:113–117.

13. Westphal M, Lamszus K: Circulating biomarkers for gliomas. Nat Rev Neurol 2015, 11:556–566.

14. Ghouzlani A, Kandoussi S, Tall M, Reddy KP, Rafii S, Badou A: Immune Checkpoint Inhibitors in Human Glioma Microenvironment. Front Immunol 2021, 12:679425.

15. Sareen H, Ma Y, Becker TM, Roberts TL, de Souza P, Powter B: Molecular Biomarkers in Glioblastoma: A Systematic Review and Meta-Analysis. Int J Mol Sci 2022, 23.

16. Bowman RL, Wang Q, Carro A, Verhaak RG, Squatrito M: GlioVis data portal for visualization and analysis of brain tumor expression datasets. Neuro Oncol 2017, 19:139–141.

17. Zhou Y, Zhou B, Pache L, Chang M, Khodabakhshi AH, Tanaseichuk O, Benner C, Chanda SK: Metascape provides a biologist-oriented resource for the analysis of systems-level datasets. Nat Commun 2019, 10:1523.

18. Subramanian A, Tamayo P, Mootha VK, Mukherjee S, Ebert BL, Gillette MA, Paulovich A, Pomeroy SL, Golub TR, Lander ES, Mesirov JP: Gene set enrichment analysis: a knowledge-based approach for interpreting genome-wide expression profiles. Proc Natl Acad Sci U S A 2005, 102:15545–15550.

19. McKinnon C, Nandhabalan M, Murray SA, Plaha P: Glioblastoma: clinical presentation, diagnosis, and management. BMJ 2021, 374:n1560.

20. Xue WC, Khoo US, Ngan HY, Chan KY, Chiu PM, Tsao SW, Cheung AN: Minichromosome maintenance protein 7 expression in gestational trophoblastic disease: correlation with Ki67, PCNA and clinicopathological parameters. Histopathology 2003, 43:485–490.

21. Walker JA, McKenzie ANJ: T(H)2 cell development and function. Nat Rev Immunol 2018, 18:121–133.

22. Li Z, Zhang Y, Sun B: Current understanding of Th2 cell differentiation and function. Protein Cell 2011, 2:604–611.

23. Li L, Yu R, Cai T, Chen Z, Lan M, Zou T, Wang B, Wang Q, Zhao Y, Cai Y: Effects of immune cells and cytokines on inflammation and immunosuppression in the tumor microenvironment. Int Immunopharmacol 2020, 88:106939.

24. Khan F, Pang L, Dunterman M, Lesniak MS, Heimberger AB, Chen P: Macrophages and microglia in glioblastoma: heterogeneity, plasticity, and therapy. J Clin Invest 2023, 133.

25. Lim M, Xia Y, Bettegowda C, Weller M: Current state of immunotherapy for glioblastoma. Nat Rev Clin Oncol 2018, 15:422–442.

26. Wang Q, He Y, Li W, Xu X, Hu Q, Bian Z, Xu A, Tu H, Wu M, Wu X: Soluble Immune Checkpoint-Related Proteins in Blood Are Associated With Invasion and Progression in Non-Small Cell Lung Cancer. Front Immunol 2022, 13:887916.

27. Kessenbrock K, Plaks V, Werb Z: Matrix metalloproteinases: regulators of the tumor microenvironment. Cell 2010, 141:52–67.

28. Reifenberger G, Wirsching HG, Knobbe-Thomsen CB, Weller M: Advances in the molecular genetics of gliomas - implications for classification and therapy. Nat Rev Clin Oncol 2017, 14:434–452.

29. Sturm D, Pfister SM, Jones DTW: Pediatric Gliomas: Current Concepts on Diagnosis, Biology, and Clinical Management. J Clin Oncol 2017, 35:2370–2377.

30. Lauko A, Lo A, Ahluwalia MS, Lathia JD: Cancer cell heterogeneity & plasticity in glioblastoma and brain tumors. Semin Cancer Biol 2022, 82:162–175.

31. Bikfalvi A, da Costa CA, Avril T, Barnier JV, Bauchet L, Brisson L, Cartron PF, Castel H, Chevet E, Chneiweiss H, et al: Challenges in glioblastoma research: focus on the tumor microenvironment. Trends Cancer 2023, 9:9–27.

32. Winkler F, Venkatesh HS, Amit M, Batchelor T, Demir IE, Deneen B, Gutmann DH, Hervey-Jumper S, Kuner T, Mabbott D, et al: Cancer neuroscience: State of the field, emerging directions. Cell 2023, 186:1689–1707.

33. Vollmann-Zwerenz A, Leidgens V, Feliciello G, Klein CA, Hau P: Tumor Cell Invasion in Glioblastoma. Int J Mol Sci 2020, 21.

34. Ludwig K, Kornblum HI: Molecular markers in glioma. J Neurooncol 2017, 134:505–512.

35. Molinaro AM, Taylor JW, Wiencke JK, Wrensch MR: Genetic and molecular epidemiology of adult diffuse glioma. Nat Rev Neurol 2019, 15:405–417.

36. Tamai S, Ichinose T, Nakada M: Liquid biomarkers in glioma. Brain Tumor Pathol 2023, 40:66–77.

37. Peng Z, Liu C, Wu M: New insights into long noncoding RNAs and their roles in glioma. Mol Cancer 2018, 17:61.

38. Cheng J, Meng J, Zhu L, Peng Y: Exosomal noncoding RNAs in Glioma: biological functions and potential clinical applications. Mol Cancer 2020, 19:66.

39. Xiao M, Wang X, Yan M, Chen W: A systematic evaluation for the potential translation of CD166-related expression as a cancer biomarker. Expert Rev Mol Diagn 2016, 16:925–932.

40. Jin S, Ye Q, Hong Y, Dai W, Zhang C, Liu W, Guo Y, Zhu D, Zhang Z, Chen S, et al: A systematic evaluation of stool DNA preparation protocols for colorectal cancer screening via analysis of DNA methylation biomarkers. Clin Chem Lab Med 2020, 59:91–99.

41. Guo L, Ding Z, Huang N, Huang Z, Zhang N, Xia Z: Forkhead Box M1 positively regulates UBE2C and protects glioma cells from autophagic death. Cell Cycle 2017, 16:1705–1718.

42. Alafate W, Zuo J, Deng Z, Guo X, Wu W, Zhang W, Xie W, Wang M, Wang J: Combined elevation of AURKB and UBE2C predicts severe outcomes and therapy resistance in glioma. Pathol Res Pract 2019, 215:152557.

43. Ma R, Kang X, Zhang G, Fang F, Du Y, Lv H: High expression of UBE2C is associated with the aggressive progression and poor outcome of malignant glioma. Oncol Lett 2016, 11:2300–2304.

44. Zang L, Kondengaden SM, Che F, Wang L, Heng X: Potential Epigenetic-Based Therapeutic Targets for Glioma. Front Mol Neurosci 2018, 11:408.

45. Xu X, Li L, Luo L, Shu L, Si X, Chen Z, Xia W, Huang J, Liu Y, Shao A, Ke Y: Opportunities and challenges of glioma organoids. Cell Commun Signal 2021, 19:102.

46. Chen R, Smith-Cohn M, Cohen AL, Colman H: Glioma Subclassifications and Their Clinical Significance. Neurotherapeutics 2017, 14:284–297.

47. Nikiforova MN, Hamilton RL: Molecular diagnostics of gliomas. Arch Pathol Lab Med 2011, 135:558–568.

48. Senhaji N, Squalli Houssaini A, Lamrabet S, Louati S, Bennis S: Molecular and Circulating Biomarkers in Patients with Glioblastoma. Int J Mol Sci 2022, 23.

49. Esparvarinha M, Madadi S, Aslanian-Kalkhoran L, Nickho H, Dolati S, Pia H, Danaii S, Taghavi S, Yousefi M: Dominant immune cells in pregnancy and pregnancy complications: T helper cells (TH1/TH2, TH17/Treg cells), NK cells, MDSCs, and the immune checkpoints. Cell Biol Int 2023, 47:507–519.

50. Lee WS, Yang H, Chon HJ, Kim C: Combination of anti-angiogenic therapy and immune checkpoint blockade normalizes vascular-immune crosstalk to potentiate cancer immunity. Exp Mol Med 2020, 52:1475–1485.

51. Chen J, Liu G, Wang X, Hong H, Li T, Li L, Wang H, Xie J, Li B, Li T, et al: Glioblastoma stem cell-specific histamine secretion drives pro-angiogenic tumor microenvironment remodeling. Cell Stem Cell 2022, 29:1531–1546 e1537.

52. Dapash M, Hou D, Castro B, Lee-Chang C, Lesniak MS: The Interplay between Glioblastoma and Its Microenvironment. Cells 2021, 10.

53. Wang T, He Z, Yuan CS, Deng ZW, Li F, Chen XG, Liu Y: MMP-responsive transformation nanomaterials with IAP antagonist to boost immune checkpoint therapy. J Control Release 2022, 343:765–776.

54. Ye Y, Kuang X, Xie Z, Liang L, Zhang Z, Zhang Y, Ma F, Gao Q, Chang R, Lee HH, et al: Small-molecule MMP2/MMP9 inhibitor SB-3CT modulates tumor immune surveillance by regulating PD-L1. Genome Med 2020, 12:83.

